# Photoreceptor cell death mechanism in light damage

**DOI:** 10.1101/2024.12.20.629734

**Authors:** Manuel G. Bruera, María A. Contin

## Abstract

Light is one of the most ambient agents harmful to the retina. In Wistar rats prolonged exposure to 200 lux of LED light promotes photoreceptors cells death after 6 days. The possibility to know the cell photoreceptors death mechanism increases the likelihood to find a mixture of therapy with inhibitors or antagonists with anti-inflammatory and antioxidant effects. This study tested the cell death mechanism to elucidate the possible underlying molecular mechanisms. Adult male rats were exposed to 200 lux of LED light by different periods of constant light. The study focused on apoptosis and necroptosis pathways. Our findings reveal that necroptosis is an active pathway in retinal degeneration, occurring in both photoreceptors and glial cells, while apoptosis seem to be inactive during the studied time points.

## Introduction

Neurodegenerative diseases represent a complex category of disorders characterized by inflammation and neuronal cell death. Photoreceptor cells are highly specialized sensory neurons that can convert light into electrical signals through the process of visual phototransduction. Death of photoreceptors leads to retinal degeneration (RD) being one of the relevant causes of blindness and may be attributed to genetic defects (Wert *et al*, 2014), or environmental factors such as detrimental habits (Zarbin, 1998) and excessive light exposure promoted by light pollution (Contin *et al*, 2015). The intracellular regulation of cell death pathways may occur by apoptosis, programmed necrosis also known as necroptosis, mitochondrial, permeability transition-driven regulated cell death, pyroptosis, and ferroptosis, which play important roles in maintaining tissue homeostasis under normal and pathological conditions. A substantial body of evidence stated that photoreceptor cell death may be made by different mechanism of cell death (Lohr *et al*, 2006; Perche *et al*, 2007; Sancho-Pelluz *et al*, 2008; Wenzel *et al*, 2005). Stress-related factors are initially addressed by innate immune cells that regulate inflammation, leading to a significant upregulation of pro-inflammatory cytokines and chemokines, as well as a marked increase in microglia/macrophages, which in turn induce neuronal cell death (Murakami *et al*, 2012; Dvoriantchikova *et al*, 2022; Wallach *et al*, 2014). Therefore, focusing on the inflammatory response could be beneficial. Inflammation damage associated molecular patterns (DAMPs) generated by necrotic cells, such as interleukin-1a (IL-1a), tumor necrosis factor a (TNF-a), and high-mobility group box 1 protein (HMGB1) strongly exacerbate inflammation (Rock & Kono, 2008; Yanai *et al*, 2009; Wallach *et al*, 2014). Previous studies have shown that prolonged exposure to low light from LED devices results in photoreceptor cell death after 6–7 days. This cell death occurs via a mechanism independent of caspase-3, with glial and immune activation revealing inflammatory processes during light exposure (Bruera *et al*, 2022; Contin *et al*, 2013). In this study, we assess the molecular mechanisms of photoreceptor cell death by focusing on signaling pathways of apoptosis caspase-3 independent and necroptosis. We proposed a caspase-3-independent mechanism of photoreceptor cell death that mediates inflammatory processes, contributing to neural damage. The animals were exposed to constant low-intensity light to induce retinal damage. The results revealed that photoreceptor cell death and microglial activation were primarily mediated by a RIP1– and RIP3-dependent necroptosis mechanism. All these results suggest that necroptotic pathways may be involved in various retinal cell types, contributing to photoreceptor death and/or sustaining inflammatory responses, which drive the progression of retinal degeneration under light stress.

## Results

### Pathways Underlying Retinal Degeneration Cell Death

Apoptosis is an essential process underlying multicellular organisms as a vital component of in cell turnover, development and functioning of the immune system, hormone-dependent atrophy, and chemical-induced cell death (Elmore, 2007). Bax and Bcl-2 are the major members of the Bcl-2 family whose potential roles are involved in several cell deaths. Bax promotes cell death through permeabilization of mitochondrial outer membrane in response to different cellular stresses and Bcl-2 prevents apoptosis by inhibiting the activity of Bax (Mohan *et al*, 2012; Maes *et al*, 2017). Previously, we demonstrated that photoreceptor cell death following light injury does not activate caspase-3, suggesting a caspase-3-independent or non-apoptotic mechanism as the major pathway of light-induced damage (Contin *et al*, 2013). To further investigate whether apoptotic mechanisms are involved in light-induced damage, retina of rats exposed to LED devices at 200 lux for 2, 4 and 6 days were processed to isolate total RNA and protein as mentioned in methods. Real-time RT-PCR analysis demonstrated that light treatment significantly enhanced BCL-2 expression following LL2, with a more pronounced increase observed at LL6 (∼1.5 fold), however, no significant differences in BAX expression were observed at any time point studied Fig. 1A. The Bax/Bcl-2 RNAm expression ratio shows significantly decreased levels from retinas light exposed (P≥0.05) compared with controls in LDR. Western blot analysis shows results in correlation with the expression of the messengers (Fig. 1B.); Bcl-2 increases under light treatment, however BAX did not show changes in the protein expression.

The cell fate decision to suffer apoptosis is influenced primarily by the activity of caspase-8 or X-linked inhibitors of apoptosis proteins activation, called *cellular inhibitor of apoptosis proteins* (CIAPs: cIAP1, cIAP2 or XIAP) (Weinlich *et al*, 2017). cIAPs governs RIPK1 ubiquitinylation and degradation (Feoktistova *et al*, 2011; Tenev *et al*, 2011) whereas caspase-8 mediates proteolytic cleavage of RIPK1 and RIPK3 (necroptosis proteins) (Lin *et al*, 1999; Feng *et al*, 2007). Caspase-8 or cIAP deficiency favors necroptosis by, respectively, removing proteolytic cleavage of RIPK1/RIPK3 or ubiquitinylation of RIPK1 (Feng *et al*, 2007). Curiously, we found that the expression of caspase-8 did not show any changes in response to light stimuli; however, and XIAP and cIAP1 showed decreased RNAm level with a significance at LL6 and LL4 respectively (Fig. 2) indicating the regulation a favor of non-apoptotic mechanism.

Cell death induction, tightly regulated by antiapoptotic proteins such as cellular FLICE-like inhibitory protein (cFLIP), a caspase inhibitor, which induce the formation of the death receptor-independent apoptotic platform known as the Ripoptosome and is also involved in the non-apoptotic cell death pathway, necroptosis (Budd *et al*, 2006). The isoform cFLIP_L_ prevents Ripoptosome formation, whereas the isoform cFLIP_S_ promotes Ripoptosome assembly (Kavuri *et al*, 2011). As is shown in Fig. 2 the isoform cFLIP_L_ showed diminutions in the RNAm expression with significance at LL6. However, the short isoform, cFLIP_S,_ involved in caspase 8 inhibition, showed increasing expression levels from LL2 indicating the non-activation or inhibition of apoptosis pathways in retinas exposed to LED light. All these results strongly suggest that apoptosis may not be the principal pathway involved in photoreceptor cell death.

### Necroptosis as a Key Pathway in Light-Induced Retinal Degeneration

*R*eceptor Interacting Protein Kinase 1 (RIPK1) and 3(RIPK3) are part of a Serine/Threonine kinases family which play a role in necroptosis through homo-or heterotypic interactions domain (Ermine et al, 2022). Phosphorylated RIP3 recruits and phosphorylates necroptotic executor Mixed Lineage Kinase Domain Like pseudo-kinase (MLKL), which translocates and oligomerizes at the plasma membrane, inducing rupture (Holler *et al*, 2000; Ermine *et al*, 2022). To analyze if necroptosis is a mechanism involved in RD by light, we analyzed the RIPK1 and RIPK3. The RNA and protein of rat retinas exposed to 2, 4 and 6 days of light (LL) were isolated as mentioned in methods. Real-time RT-PCR analysis showed non-significant increased levels of RIPK1 (Fig. 3A). However, light treatment increases the expression of RNAm of RIPK3 with significant value at LL6. The MLKL RNAm analysis showed no significatively increasing levels in any time of LL studied; nonetheless, the immunohistochemistry analysis of phosphorylated protein showed positive immunostaining in retinas after 2 days of light treatment (Fig. 3B and B). The positive labeling is visualized for different retinal cells, including the outer segments of the photoreceptors (Figure 3C). The analysis with IB4, an isolectin marker of microglia retinal cells, showed positive co-staining at LL4 and LL6 indicating an increase in p-MLKL expression in glial cells after light injury (Fig. 3C and 4B anc C). Finally, the analysis of TNF, TNFR1 and TLR4 RNAm expression showed increased levels of RNAm expression at LL4 and LL6 in the isoform TNFR1 and TNFR2 and increased levels in TLR4 at LL2 (Fig. 4A).

## Discussion

Even though the retina is responsible for perceiving light for vision, excessive exposure makes it vulnerable, especially to the photosensitive cells (Williams & Howell, 1983; Gorn & Kuwabara, 1967; Williams & Penn, 1985; Organisciak *et al*, 1989a, 1989b) (Bicknell et al., 2002; Contin et al., 2013; Hajkova et al., 2010; Nakamura et al., 2018; Organisciak, Wang, et al., 1989a, 1989b) (Contin et al., 2015; Marc et al., 2008; Sang et al., 2011). Light-emitting diodes (LEDs) are the major domestic and public light source and are in computers, tablets, cell phones and game consoles, promoting high artificial exposure of the visual system. Today, we are exposed to more light than we were twenty years ago, yet there is a lack of political attention on educating the public about the effects of new lighting technologies, particularly regarding excessive nighttime exposure. It is important to recognize that the potential harm to the visual system should be thoroughly evaluated. Exposure to artificial light is an environmental factor that can affect physiological processes and potentially contribute to retinal aging. This can lead to age-related macular degeneration or even hasten the progression of genetic diseases like retinitis pigmentosa (Marquioni-Ramella & Suburo, 2015)

In previous studies, we developed a retinal degeneration (RD) model by exposing adult Wistar rats to low-intensity white LED light (200 lux) for periods of 1 to 8 days (LL1-LL8). We observed significant photoreceptor cell death after 5 to 6 days of continuous exposure (LL5-LL6) (Contin et al., 2013). Under light/dark exposure conditions, the animals showed early retinal injury after 7 days of light/dark cycles (LD7). This suggests that, even with intermittent periods of darkness, LED exposure may result in visual damage (Benedetto *et al*, 2023).

We further examined the role of caspase-3 in cell death and found no evidence of its activation at any of the time points studied (Contin et al., 2013). Understanding the molecular pathways involved in cell death is crucial for developing potential therapies to prevent or mitigate retinal degeneration. Cell death pathways are regulated at multiple levels, with organelles such as mitochondria being key initiation points (Kelly & Strasser, 2011). Caspases are a family of cytosolic cysteine proteases that cleave their substrates after an aspartic acid residue. Initiator caspases include caspases 2, 8, 9, and 10, while effector caspases include caspases 3, 6, and 7 (Degterev *et al*, 2003). There are seven human inhibitor of apoptosis proteins (IAPs), including X-linked IAP (XIAP), c-IAP1, c-IAP2, NAIP, Surviving, BRUCE, and ML-IAP (Deveraux *et al*, 1997; Huang *et al*, 2001). XIAP is a crucial regulator of caspases and is subject to complex negative regulations. It plays multifunctional roles in regulating apoptosis and various cell signaling pathways (Holcik & Korneluk, 2001; Holcik *et al*, 2001). XIAP is the most potent and versatile member of apoptotic regulators, controlling caspases 3, 7, and 9. Caspase-independent mechanisms also play a role in the intrinsic death pathway, downstream of Bax and Bak. These proteins maintain mitochondrial integrity and facilitate the release of pro-apoptotic molecules from mitochondria (Hevler *et al*, 2021). Inhibitors of apoptosis proteins (IAPs) add an additional regulatory layer to intrinsic apoptotic pathways to prevent unnecessary caspase activation (Deveraux *et al*, 1997). In this study, we demonstrate elevated levels of Bcl-2, accompanied by a corresponding decrease in the Bax/Bcl-2 ratio, in the retinas of animals exposed to LED light. Additionally, the expression of the inhibitors cIAP and XIAP was also reduced. These findings suggest that apoptosis is, at most, a non-dominant mechanism involved. Thus, these findings suggest that the molecular mechanisms underlying RD do not involve cell death through either caspase-dependent or caspase-independent pathways.

Necroptosis, on the other hand, is involved in a variety of cell death by diseases such as atherosclerosis, pancreatitis, stroke, myocardial infarction, ischemia-reperfusion injury and inflammation. Is a regulated cell death with intracellular core components of the necroptotic machinery, RIPK3 and MLKL controlled principally by receptor-interacting protein kinase 1 (RIPK1) and RIPK3 which activates mixed lineage kinase domain-like pseudo kinase (MLKL) (Cho *et al*, 2011). Moreover, the caspase inhibitor cFLIP, induce the formation of Ripoptosome (a receptor-independent apoptotic platform) involved in the necroptosis (Budd *et al*, 2006) promoting the necrosome, a complex of the three proteins RIPK1-RIPK3-MLKLwhich may be stimulated by several different factors such as pathogen infection, lipopolysaccharides, TNF superfamily, interferon-γ, and various drugs (Chen *et al*, 2014). TNF-α is the receptor that mediates signaling pathway, cFLIP regulates both extrinsic apoptosis pathway and necroptosis pathway.

In this work, the analysis of the regulated cell death, necroptosis, showed increased levels of RIPK3 RNAm from LL2 to LL6 with statistical significance at LL6. The analysis of RNAm expression showed no changes in the levels of the large form, cFLIP_L_ being significant only after LL6, however, there are elevated levels of short form (cFLIPs) at LL2 indicate that the Caspase-8-cFLIPS heterodimer lacks the proteolytic activity necessary not only for full Caspase-8 activation but also for RIP1 degradation. Consequently, RIP1 accumulates within the ripoptosome and triggers necroptosis via RIP3 in a RIP1 kinase-dependent manner (Feoktistova et al, 2011a). Necroptosis is initiated upon activation of the receptor interacting serine-threonine kinases RIPK1 and RIPK3 after TNF receptor 1 (TNFR1) or Toll-like receptor (TLR) stimulation. RIPK3 phosphorylates the pseudo-kinase mixed-lineage kinase domain-like (MLKL), which undergoes conformational modifications causing it to translocate and insert into the cell membrane. At the cell membrane, MLKL induces membrane rupture (possibly via pore formation) (Kavuri *et al*, 2011); so MLKL plays a key role in induction of necroptosis. MLKL acts in two ways: (1) either acts as platform in plasma membrane for recruitment of Na+ ion or Ca++ channels or (2) promotes the pore formation in plasma membrane by interacting with amino terminal of phosphotidyl inositol phosphate (Dhuriya & Sharma, 2018; Tang *et al*, 2009). Here we demonstrated that the RNAm expression of MLKL did not show changes in any times of LL studied; however, the IHC showed the protein expression in both photoreceptor and glial cells. Glial cells serve as defense, playing a pivotal role in balancing both cytotoxic and cytoprotective effects at the intersection of homeostasis and disease (Rodríguez-Gómez *et al*, 2020). Persistent activation of microglia can trigger necroptosis, a type of necrosis induced by the activation of receptor-interacting protein kinase 1 (RIPK1)/RIPK3 (Weinlich *et al*, 2017; Galluzzi *et al*, 2018), and necroptosis in turn induces secretion of proinflammatory DAMPs and mediates chronic inflammation (Rodríguez-Gómez et al., 2020). It has been shown that microglial necroptosis executed by RIPK1/3 contributes to neuronal damage as well as brain regeneration (Welser *et al*, 2010; Ofengeim *et al*, 2017; He *et al*, 2021). Previously, we demonstrated an increased number of Iba1-positive cells in the outer nuclear layer, which exhibited the characteristic morphology of activated microglia. In retinal inflammation, it has been suggested that microglia trigger necroptosis, which exacerbate retinal neural damage and degeneration. (Huang *et al*, 2018). Furthermore, we observed increased levels of immune mediators TNF, IL-6, and the chemokines CX3CR1 and CCL2, all indicative of glial activation, which appeared after 5-6 days of constant light exposure. By this time, the number of photoreceptor cells had already significantly decreased, suggesting that glial and immune activation occurs secondary to neurodegeneration. This implies that photoreceptor cell death is an early event that may occur independently or concurrently with glial-derived immune responses. Oxidative stress could be linked to glial activation following prolonged light exposure (Bruera et al, 2022). ROS may activate NF-κB which is a vital transcription factor included in immune reaction and inflammation. Activated NF-κB translocate from the cytoplasm to inside the nucleus where it can bind with the promoter regions enhancing transcription of several genes, including, Cox2 and tumor necrosis factor-alpha (TNF-α) and thus, promoting inflammation (Liu *et al*, 2017). When the phase of acute inflammation is induced it is uncontrolled, and excessive inflammation may become chronic and harm several cells of the retina. So, the results showed that necroptosis mechanisms play different roles depending on the retinal cell type of activation.

Studies conducted of retinal transplant by Maidana and col. demonstrated photoreceptor survival in RIP3 kinase deficiency in both donor and recipient tissue. The deletion of RIPK3 in the peripheral immune cells of recipient animals significantly protected photoreceptor graft survival. These effects were also replicated by pharmacological inhibition of the RIPK1-RIPK3 axis, highlighting the critical role of RIPK in retinal degeneration through both local and systemic immune cell interactions (Maidana *et al*, 2023) the authors conclude that targeting the RIPK3 pathway can help promote cell survival. A combination of immunomodulatory and neuroprotective strategies targeting necroptotic pathways may contribute to restoring photoreceptor loss in retinal degenerations.

Based on these findings and the results presented here, we conclude that necroptosis is an active pathway in retinal degeneration promoted by exposure to LED light. This pathway is expressed in both photoreceptors and glial cells, while apoptosis is not activated during the studied time points.

Immunomodulatory and neuroprotective strategies targeting the necroptosis pathway could support therapies aimed at preventing retinal degeneration.

## Material and Methods

### Animals

According to the 3Rs principles for ethical use of animals in scientific research, all efforts were made to minimize both animal numbers and their suffering. Male albino Wistar rats (12–15 weeks), inbreed in our laboratory for 5 years, were maintained on 12:12 h light-dark cycles with lights on (less than 50 lux of white fluorescent lamp) from zeitgeber time (ZT) 0 to 12 from birth until the day of the experiment. Food and water were available *ad libitum*.

### Retinal Light Damage

Retinal degeneration was induced as described by Contín et al (Contín et al., 2013). Briefly, animals were exposed to constant light in boxes equipped with LED devices (EVERLIGHT Electronic Co., Ltd. T-13/4 3294-15/T2C9-1HMB, color temperature of 5,500 K) in the inner upper surface and temperature-controlled at 24 ± 1°C. At rat eye level, 200 lux were measured with a light meter (model 401036; Extech Instruments Corp., Waltham, MA, USA). After light stimulation the animals were sacrificed in a CO_2_ chamber at ZT6.

### Light Exposure Protocol

Animals were exposed to constant light stimulation for 2, 4, and 6 days (LL2-LL6). Rats exposed to fluorescent light at 50 lux on 12:12 h light-dark cycles (LDR) were used as controls of standard housing conditions.

### RNA isolation and cDNA synthesis

Total RNA was extracted from rat retinas using TRIzol® RNA extraction reagent (Invitrogen, Carlsbad, CA), according to the manufacturer’s instructions. The isolated RNA was quantified using an Epoch Microplate Reader (BioTek Instruments, Winooski, VT, USA). For complementary DNA (cDNA) synthesis, 2 µg of RNA was treated with DNase I (Thermo Scientific, USA) to remove possible contamination with genomic DNA. The product was incubated with a mix of random hexamer and Oligo-dT primers (Biodynamics), deoxynucleotides and the reverse transcriptase M-MLV (Promega, Madison, WI, USA), in RNAse-free conditions. Reverse transcription was performed following the manufacturer’s specifications, employing a thermocycler Mastercycler gradient (Eppendorf, Hamburg, Germany).

### Real time RT-PCR

cDNA (120 ng) was amplified in 15 μl reaction mixture consisting of 7.5 ul of 2x SYBR Green PCR Master Mix (Life Technologies, USA), 0,75 μl 10 μM primer mixture and 0,75 μl of Nuclease-Free Water with an CFX96 Touch Real-Time PCR Detection System (Bio-Rad, USA). The parameters used for PCR were as follows: 95°C for 5 min (1 time); 95°C for 30 sec, 60°C for 30 sec, and 72°C for 30 sec (40 times) and 95°C for 60 sec (1 time). Samples were subjected to a melting-curve analysis to confirm the amplification specificity. Semi-quantification was performed by the method of ΔΔCt. The fold change in each target gene relative to the □-Actin endogenous control gene was determined by: fold change = 2-Δ(ΔCt) where ΔCt = Ct(target) – Ct(□-Actin) and Δ(ΔCt) = ΔCt(LL) – ΔCt(LDR). Real time RT-PCR were run separately for each animal in triplicate. The primers used for real time RT-PCR are provided in Table 1.

### Immunoblot

Homogenates of whole retina suspended in 250 μl of RIPA buffer [50 mM Tris–HCl pH 8.0; 150 mM NaCl; Igepal (NP-40), 1% (v/v); sodium deoxycholate, 0.5% (w/v); and SDS, 0.1% (w/v)] supplemented with protease and phosphatase inhibitors were lysed through repeated cycles of ultrasonication. Total protein content was determined using the Pierce BCA Protein Assay Kit (Thermo Scientific, #23227). Lysates were then mixed with sample buffer [62.5 mM Tris–HCl pH 6.8; SDS, 2% (p/v), glycerol, 10% (v/v); 2-mercapto-ethanol, 5% (v/v); bromophenol blue, 0.002% (w/v)] and heated at 90°C for 5 minutes. Proteins (70 μg per lane) and molecular weight marker [5 μl of Precision Plus Protein All Blue Standards, range 10–250 kDa, from BIORAD (#1610373)] were separated by SDS-gel electrophoresis on 12% polyacrylamide gels. The separated proteins were transferred onto nitrocellulose membranes, blocked for 1 hour at room temperature (RT) with blocking buffer [3% (w/v) BSA in washing buffer containing 0.1% (v/v) Tween-20 in Tris-buffered saline (TBS, pH 7.4)], and subsequently incubated overnight at 4°C with primary antibodies diluted in blocking buffer. The following primary antibodies were used: rabbit anti-Bcl-2 (1:1000 dilution, Cell Signaling Technology, #2870), rabbit anti-Bax (1:500 dilution, Cell Signaling Technology, #14796), and mouse anti-α-Tubulin (1:2000 dilution, Sigma Aldrich, #T9026). After washing with washing buffer, the membranes were incubated for 1 hour at RT with the corresponding secondary antibodies (1:10000 dilution, Goat anti-rabbit IRDye ® 700CW or Goat anti-mouse IRDye ® 800CW, Odyssey LI-COR) in TBS, followed by three washes (5 min each) with washing buffer. Membranes were scanned using an Odyssey IR Imager (LI-COR Biosciences), and protein bands were quantified by densitometry using the FIJI/ImageJ software (NIH).

### Intravitreal injection of Propidium Iodide

After light exposure, the rats were anesthetized via intraperitoneal injection of a combination of xylazine hydrochloride (2 mg/kg) and ketamine hydrochloride (150 mg/kg). Local anesthesia was administered using proparacaine hydrochloride (0.5%; Alcon Laboratories). Intravitreal injection of propidium iodide (Sigma Aldrich) was performed under a surgical microscope using a 30-gauge needle attached to a calibrated Hamilton microsyringe. Propidium iodide was delivered at a concentration of 1 μg per eye in a volume of 8 µL of phosphate-buffered saline (PBS, pH 7.4). Following intravitreal injection, the animals were kept in darkness for 3 hours before being sacrificed for tissue collection.

### Immunohistochemistry

After light treatment, the rat eyes were immediately enucleated and fixed overnight (ON) at 4°C in 4% (w/v) paraformaldehyde (PFA) in PBS. Following fixation, the eyes were washed in PBS and sequentially cryoprotected in 15%, 20%, and 30% sucrose solutions. The cornea, lens, and vitreous body were carefully removed, and the eyecups were embedded in optimal cutting temperature (OCT) compound (Tissue-Tek®; Sakura) and frozen. Retinal sections, 12 μm thick, were obtained along the horizontal meridian (nasal-temporal) using a cryostat (HM525 NX-Thermo Scientific). The sections were washed in PBS and permeabilized in 0.25% (v/v) Triton X-100 (Sigma Chemical Co., St. Louis, MO, USA) in PBS for 30 minutes at RT. Subsequently, they were blocked in blocking buffer [PBS containing 0.25% Triton X-100, 3% (w/v) BSA, 1% (w/v) glycine, and 0.02% (w/v) sodium azide] for 1 hour and 30 minutes at RT with continuous gentle shaking. After that, sections were incubated with rabbit anti-Phospho-MLKL (dilution 1:500, Cell Signaling Technology, #37333) diluted in blocking buffer, ON at 4°C in a humidified chamber. Samples were rinsed three times for 5 min each in PBST [PBS containing 0.05% (v/v) Triton X-100] and then incubated for 1 hour at RT with donkey anti-rabbit IgG conjugated to Alexa Fluor 488 (711-545-152; Jackson ImmunoResearch), diluted 1:1000 in PBS containing 0.25% (v/v) Triton X-100, 1% (w/v) BSA, and 0.02% (w/v) sodium azide. Cell nuclei were labeled with 3 μM DAPI. Finally, samples were washed three times in PBST and mounted in Mowiol (Sigma-Aldrich Co., St. Louis, MO, USA). Microglial cells were labeled with biotin-conjugated isolectin B4 (IB4; 1:150 dilution, Sigma-Aldrich, L2140), followed by incubation with streptavidin conjugated to Alexa Fluor 555 (1:1000 dilution, Invitrogen, S32355). Images were acquired using a confocal microscope (Olympus FV1200, Japan).

#### Retinal Flat Mount

After fixation of the eyes in PFA and washing with PBS, the retina was dissected and separated from the choroid and pigment epithelium under a surgical microscope. Retinas were transferred to a 24-well plate, washed three times for 5 min each with PBS, and blocked for 3 hours at RT in blocking solution [PBS containing 0.5% Triton X-100, 3% (w/v) BSA, 1% (w/v) glycine, and 0.02% (w/v) sodium azide] with continuous gentle shaking. Subsequently, the retinas were incubated overnight at 4°C with anti-pMLKL primary antibody and isolectin IB4, both diluted in blocking solution. The following day, the samples were washed three times for 5 minutes each with PBST and incubated with DAPI, the appropriate secondary antibody, and streptavidin conjugate for 3 hours at RT with gentle shaking. Finally, the retinas were washed with PBST, mounted photoreceptor side up on slides, and covered with Mowiol.

#### Statistical Analysis

Statistical analysis was carried out using the Infostat software (Version 2017, InfoStat Group, FCA, National University of Cordoba, Argentina). The assumptions of normality and homogeneity of the variance were proved by Shapiro-Wilks and Levene tests, respectively. When data was normally distributed was analyzed using one-way analysis of variance (ANOVA) and Bonferroni post hoc test. When the data did not comply with the assumptions of the ANOVA tests being non-normally distributed for the groups, data were analyzed using a non-parametric Kruskal-Wallis test. Data are expressed as mean ± SD. In all cases, a p value < .05 was considered statistically significant. All graphics were made using GraphPad Prism Software, version 6.01 (San Diego, CA, USA).

## Data availability section

This study includes no data deposited in external repositories.

## Acknowledgements

The authors thank TRB Argentina for their support. Express their gratitude to Dr. Cecilia Sampedro and Dr. Carlos R. Mas for their invaluable technical support in image acquisition and analysis. Special thanks are extended to Rosa Andrada for her exceptional management of the animal facility. This work was made possible by the generous support of grants from the Agencia Nacional de Promoción Científica y Técnica (PICT 2020 No. 02699), Consejo Nacional de Investigaciones Científicas y Tecnológicas de la República Argentina (CONICET PIP 2020), Secretaría de Ciencia y Tecnología de la Universidad Nacional de Córdoba (SeCyT-UNC), and the Ministry of Sciences and Technology of Córdoba.

## Author contributions

**Manuel G Buera**: Conceptualization; Conducting the experiments; Formal analysis; Validation; Investigation; Visualiza-tion; Methodology.

**Maria Ana Contin:** Conceptualization; Supervision; Funding acquisition; Writing— original draft; Project administration; Writing—review and editing.

---

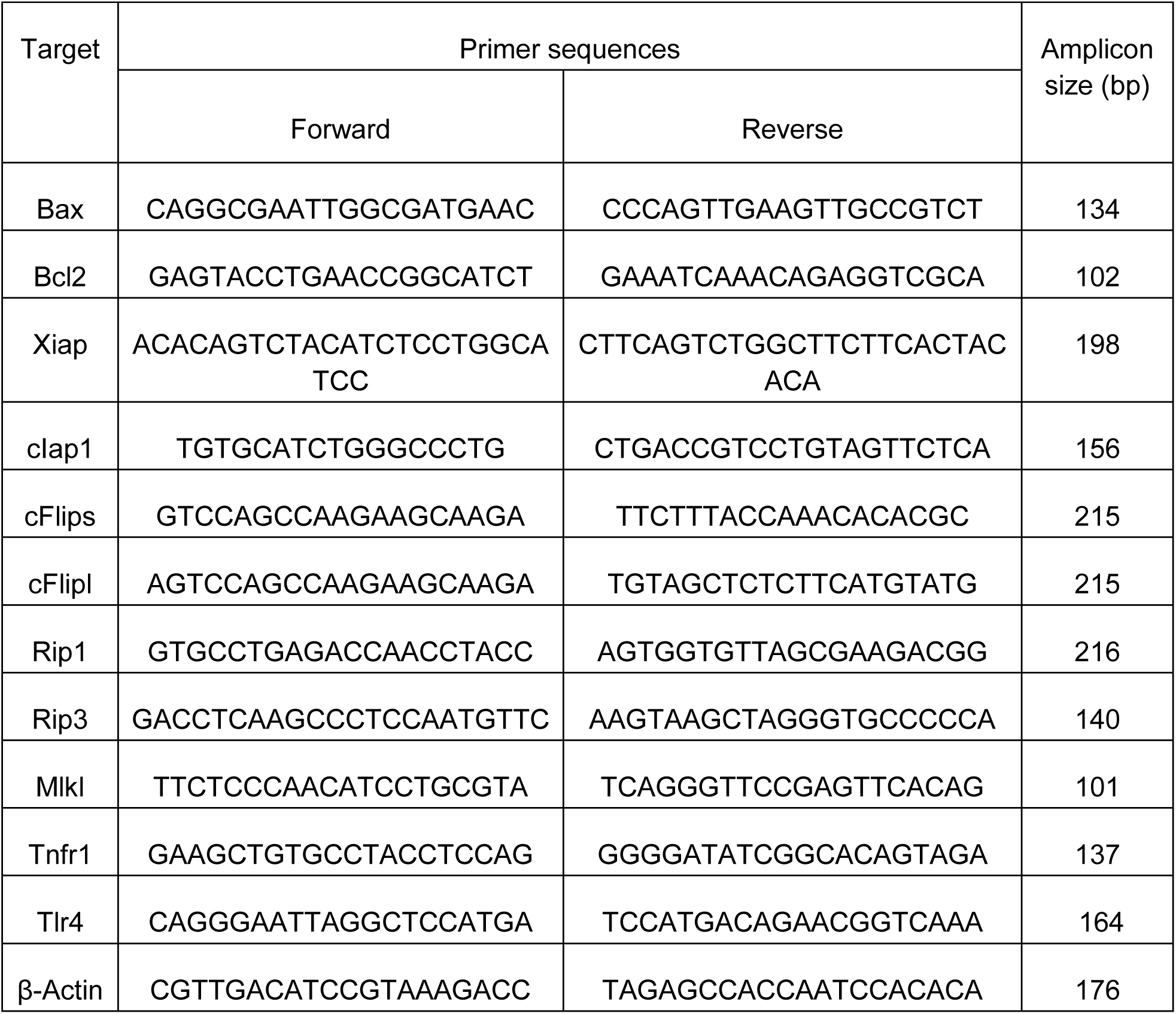
Table 1.

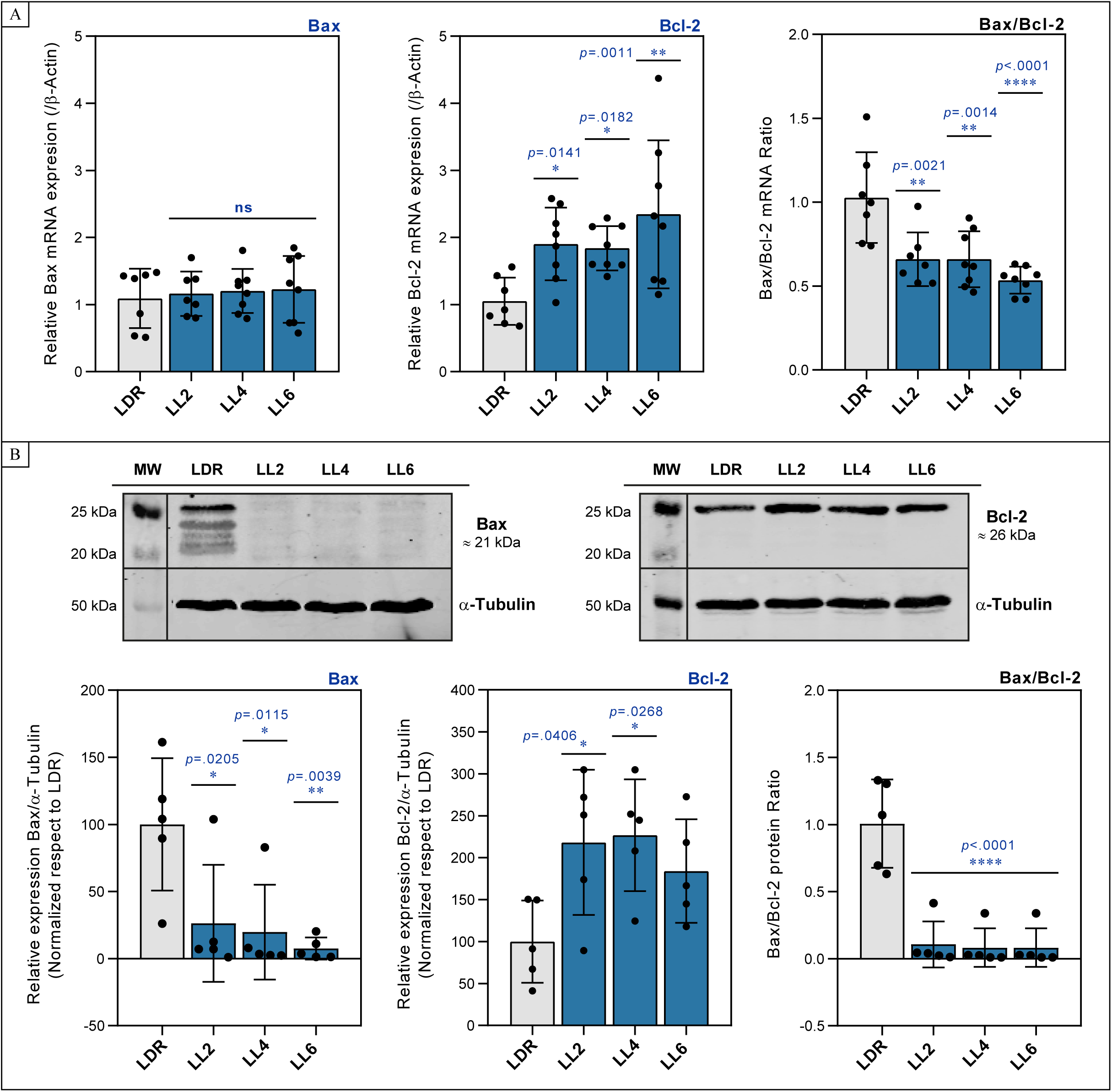

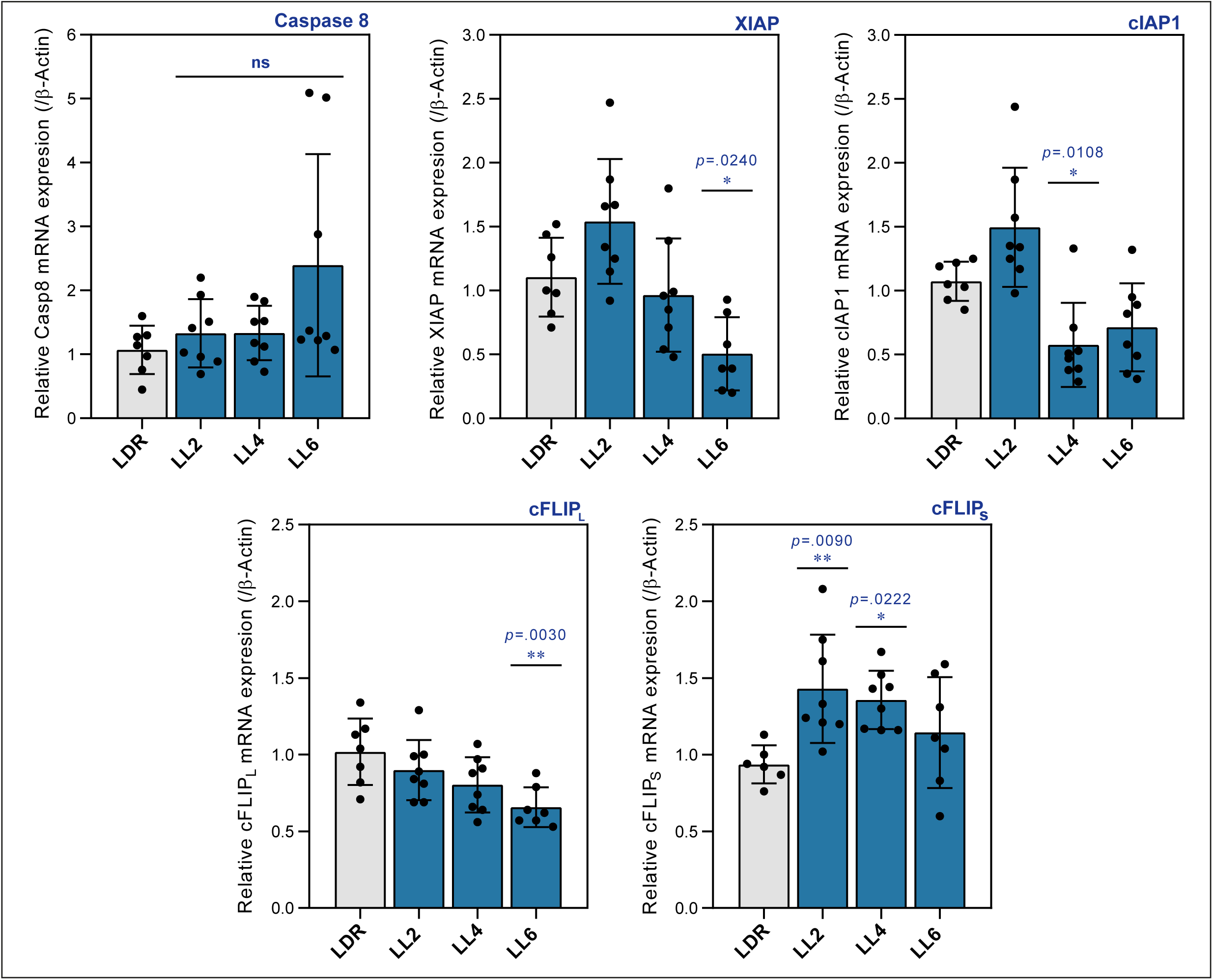

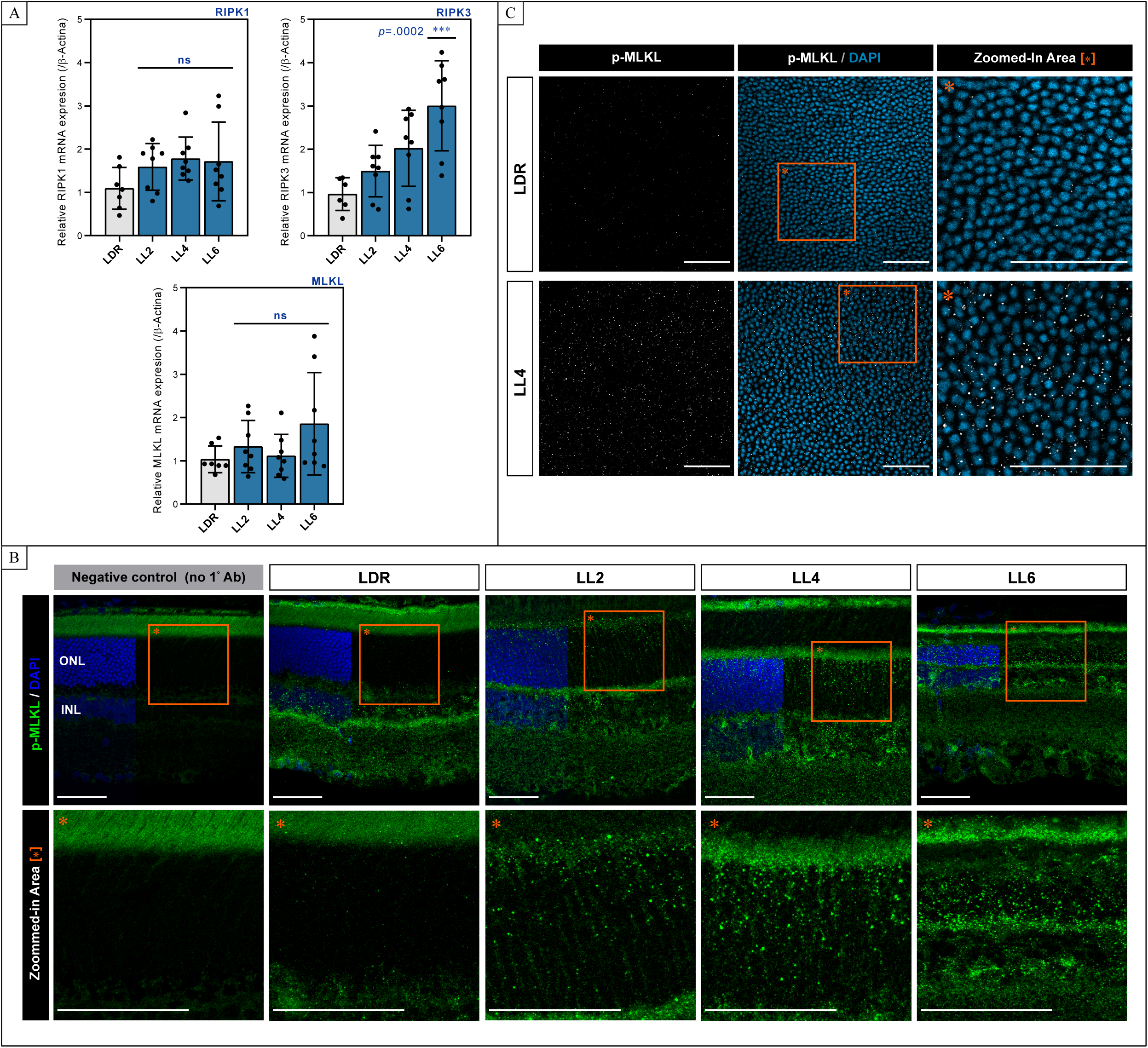

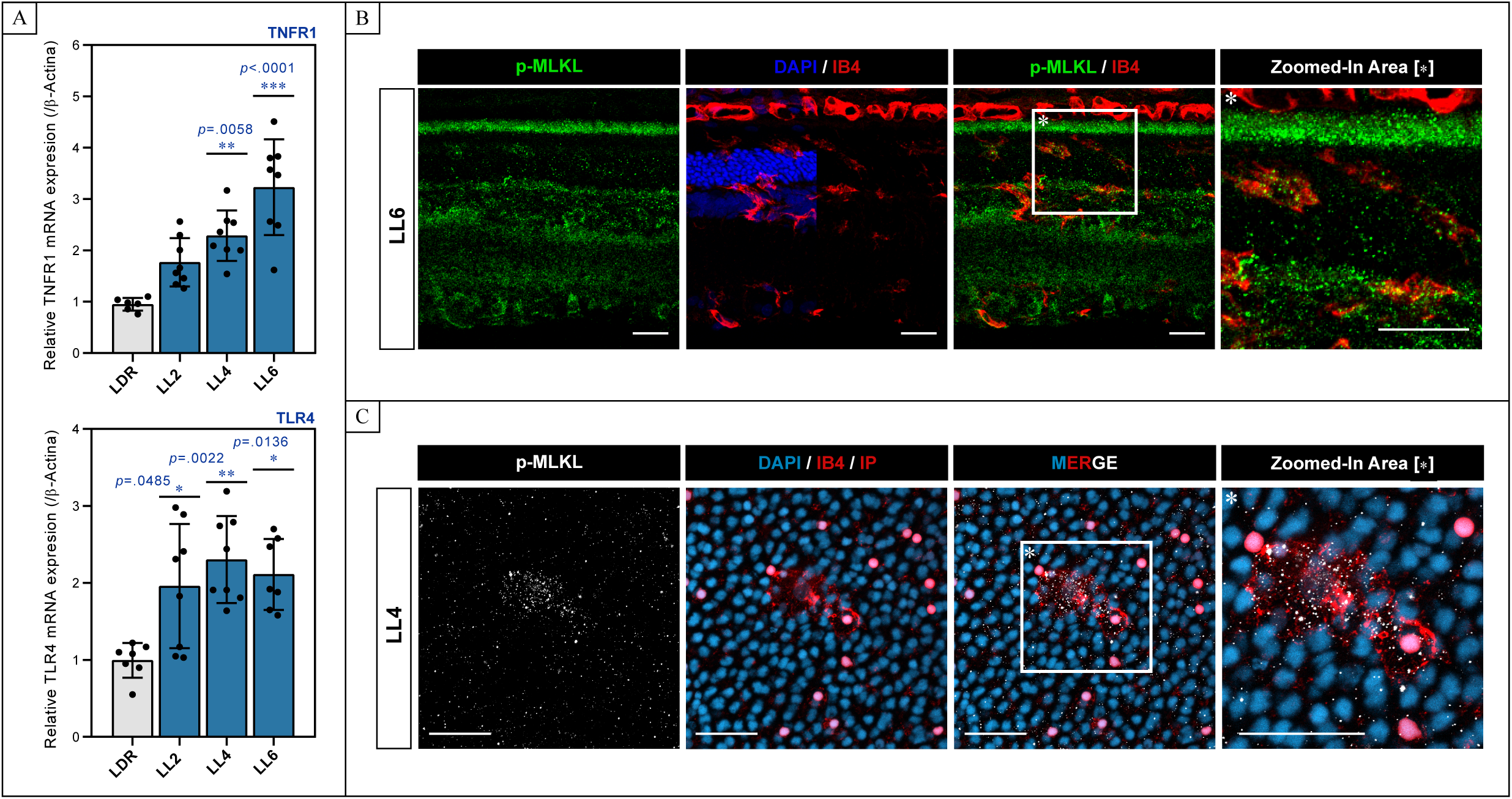

